# Computational model for migration of human osteoblasts in direct current electric field

**DOI:** 10.1101/2020.12.15.422893

**Authors:** Jonathan Edward Dawson, Tina Sellmann, Katrin Porath, Rainer Bader, Ursula van Rienen, Revathi Appali, Rüdiger Köhling

## Abstract

Under both physiological (development, regeneration) and pathological conditions (cancer metastasis), cells migrate while sensing environmental cues in the form of mechanical, chemical or electrical stimuli. Although it is known that osteoblasts respond to exogenous electric fields, the underlying mechanism of electrotactic collective movement of human osteoblasts is unclear. Theoretical approaches to study electrotactic cell migration until now mainly used reaction-diffusion models, and did not consider the effect of electric field on single-cell motility, or incorporate spatially dependent cell-to-cell interactions. Here, we present a computational model that takes into account cell interactions and describes cell migration in direct current electric field. We compare this model with *in vitro* experiments in which human primary osteoblasts are exposed to direct current electric field of varying field strength. Our results show that cell-cell interactions and fluctuations in the migration direction together lead to anode-directed collective migration of osteoblasts.

**Author summary:** Electrotactic migration of cells involves directed movement of a large number of single cells under the influence of external electric field. Influencing the migration behaviour of osteoblasts by external direct current electric field offers a promising approach towards building highly effective implants for bone regeneration. We present a computational model for electrotactic migration of osteoblasts subject to external direct current electric field. Our model considers individual cells that interact with each other and the external electric field, and, replicates the experimental observations, based on single-cell analysis, of the response of osteoblasts to electrical stimulation of varying strengths for 7 hours. Our results suggest that tracking trajectories of individual cells provide a way of determining the role of various interactions of a cell in collective migration. Our model provides a framework that links single cell response to the large scale collective dynamics.

## Introduction

The response of the cell to its sensory inputs plays a crucial role in many biological processes such as embryonic development, tissue formation/regeneration and wound healing. One of the crucial common reactions of cells is their directed motility, where cells alter their motion in response to external stimuli. Generally, such stimuli are considered to consist of chemical (chemotaxis) or mechanical (adhesion and substrate contact; haptotaxis) mechanisms, as well as of temperature gradients (thermotaxis) or electric fields (electrotaxis) [1], [2], [3, 4]. The latter, electrotaxis, also termed galvanotaxis, is increasingly studied in particular in keratinocytes and fibroblasts, since it may provide a promising strategy to foster skin wound healing [5], [6], [7], [8]. In this context, several groups aimed to clarify the nature of the electric field sensor. One possible candidate of such a sensor is the outer, negatively charged glycocalyx, which also is responsible for adhesive behaviour [9]. Other studies point to an important role of lipid rafts: their redistribution and clustering appear to be responsible for electrical field sensing in fibroblasts, mesenchymal stem cells, and adenocarcinoma cells [7], but also in corneal epithelial cells [10]. In most of these cells, the orientation seems to be cathodal (e.g. in fibroblasts and mesenchymal stem and corneal epithelial cells [7] [10]). However, this does not apply to all cell types: adenocarcinoma cells, and bone marrow mesenchymal stem cells, show the opposite orientation, i.e. anodal [7, 11]. Also the downstream signalling apparently is differential; in the various studies, Rho and PI3K [7], EGF and ERK1/2 [10], or PKG, and again PI3K (this time in dictyostelium [12], where starvation appears to initiate migratory movement [13]) were found to be involved. Interestingly, a reversal of directionality was reported for keratinocytes when inhibiting P2Y receptors [8]. We recently reported that store-operated calcium channels are pivotal for electrotaxis in human osteoblasts [14], which interestingly migrate to the anode. Thus, both electrotaxis as such, as well as the polarity, seem to be dependent on a variety of factors, such as cell type, environment, possibly age and ontogenetic stage, all of which should influence signalling pathway equipment.

One of the factors that has not found much consideration so far: *In vivo*, electrotactic cell migration involves not only singular, but many cells, for example in a tissue, which collectively respond to either endogenous or exogenous electric fields. Such an electric field-dependent collective cell migration raises the question in which way electric field on the one hand, and neighbour cell behaviour on the other (both close-range limited by finite volume, and intermediate governed by group orientational alignment) interact to generate a final migration vector. In previous modeling studies, mainly reaction-diffusion based models were used [15], [16], in some cases including interaction between electrical field and chemoattractant [17], [3]. The focus of these approaches was on cell migration mainly at the mean-field level and did not resolve the processes at the level of a single cell. Thus, cell-cell interactions as possible determining factors for cell migration direction and speed have not been modeled so far. Cell-cell communication establishes a network which gives rise to many interesting behaviours, such as non-linear collective response, as observed in quorum sensing, a type of bacterial cell-cell communication [18, 19]. Quantitative studies have shown that collective cell migration in epithelial structures is an emergent phenomenon, that cannot be explained without considering cell-cell interactions [20, 21]. A specific class of agent-based model that takes into account interactions between individual active particles during migration in continuous space are the self-propelled particle models, which were developed to understand flocking phenomena and show that under some conditions transitions can be observed where collective effects give rise to a common motility pattern [22–24]. Self-propelled particle-based models have been widely used to study collective behaviour in cell migration in tissues [25, 26]. Self-propelled voronoi model, a hybrid of self-propelled particle model and vertex model, that links active cell mechanics with cell shape and cell motility, predicts a liquid-solid transition in confluent tissues, where cell-cell interactions, among others, play a key role [27], [21, 28]. While inclusion of cell-cell interactions in models seem to be natural in the case of high-density tissue culture, where cells adhere to each other and thus exert a pulling force on the neighbouring cells, for examples in epithelial wound healing [29], the rules governing such an interaction in a system of isolated cells, such as *in vitro* cell culture, remains ambiguous. To our knowledge, to date no computational model has taken into account individual cell interactions to study the migration of cells stimulated by external electric field. Here, we present a computational model for collective dynamics of cells stimulated by direct current (DC) electric field. Our modeling framework incorporates motion of single cells, fluctuations in the direction of cell migration, cell-cell interactions, and cell-electric field interactions. By re-analysing data on individual cell basis from our recent study on osteoblast migration mechanisms in DC electric field [14], we test the hypothesis that cell-cell interactions shape the total vector.

## Results

### In vitro DC stimulation of human osteoblasts

In the experimental part of this study, we exposed human osteoblasts to DC electric fields for 7 hours (h) at different stimulation strengths and matched each of these experiments with a sham-stimulated, control group treated identically, save the DC stimulation. For the analysis of the migration behaviour, we selected adherent cells in the stimulation chambers which could be identified clearly at starting and end points of the experiment, and did not form clusters precluding the outlining of their boundaries (Fig 1 A-C). Using photographs of several fields of vision in each chamber, 1-4 cells could be traced in this way per field of vision position, totalling n=177 cells (sham stimulation), as well as n=34 cells (at 160 V/m), n=35 cells (at 300 V/m), n=26 (at 360 V/m), n=43 (at 425 V/m) and n=33 (at 436 V/m). As one can notice in the original photographs of one typical cell from the experiment using 436 V/m stimulation, the cells move (in this case anodally), and at the same time change their shape within the 7 h stimulation (Fig 1 A-C). While we did not analyse shape changes any further in this study, we took them into consideration by using centroids of the cells (blue dots in Fig 1 C) as markers to determine the net movement.

**Fig 1.**
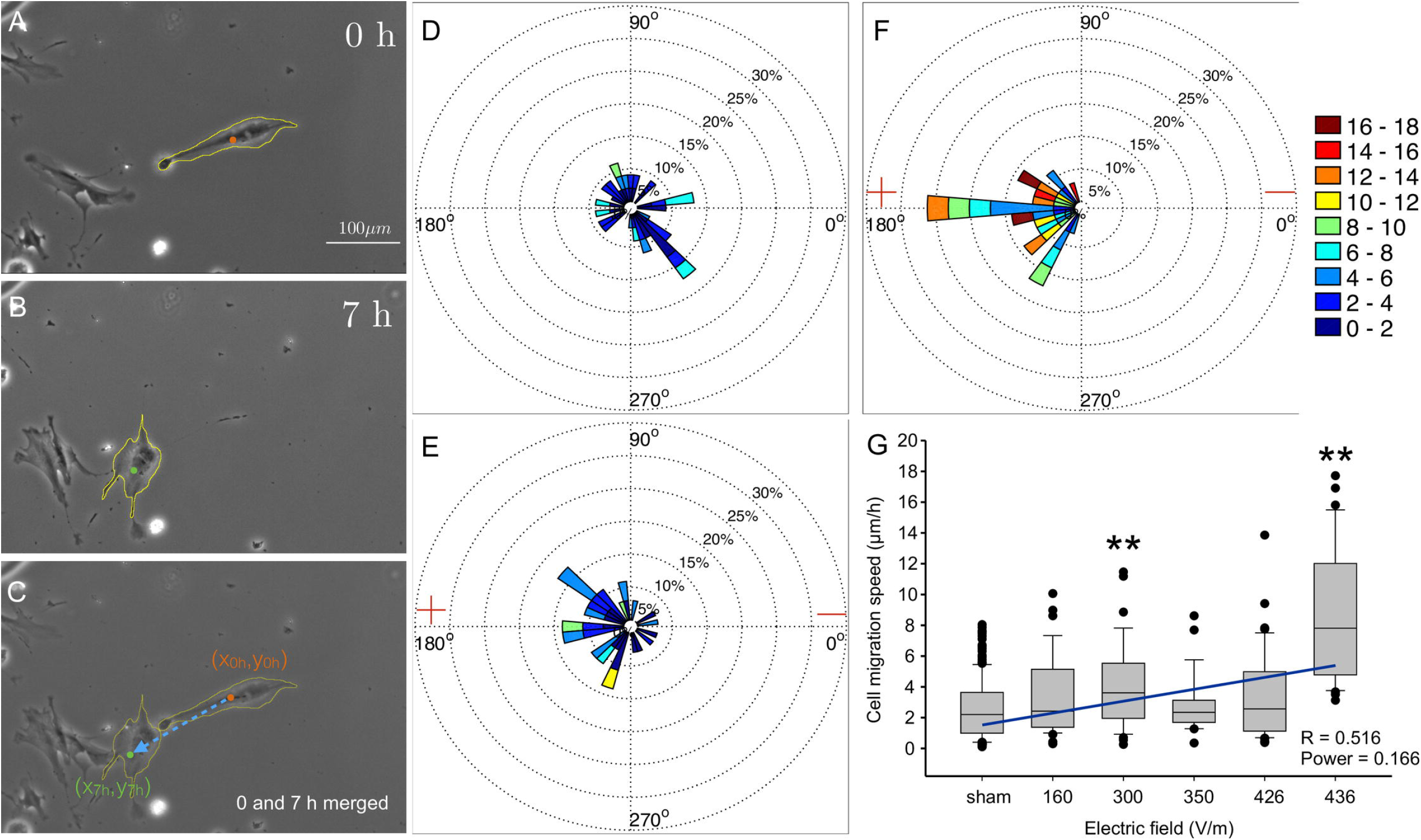
Single-cell analysis of migration of osteoblasts in a DC electric field. (A-C) Photomicrographs of osteoblasts in stimulating chamber. Cell boundaries were manually outlined as depicted (yellow coastline). (A) Time point before stimulation. (B) Position of the same cells after 7h DC-stimulation (436 V/m). (C) Overlay of A and B. Blue arrow shows the displacement of the centroid of the same cell before and after 7 h DC-stimulation. (D-F) Polar plots showing the velocity of cell migration in the cases of sham D, 160 V/m, E, and 436 V F DC stimulation. Each polar plot has been divided into 36 sectors, and data of cells migrating within each 10° sector are cumulated. Speed range is colour coded (in *μ*m/h) as shown in the colourbar. The relative sector lengths denote the percentage of cells migrating at a certain speed range. (G) Box and whisker plot of medians (horizontal lines) of cell migration speed vs. electric field strength. Whiskers denote 25-75 percentiles of data distribution. Dots show data lying outside these percentiles. Numbers of cells for each experiment are: 177, 34, 35, 26, 43, 33 for sham, and 160, 300, 360, 426 and 436 V/m, respectively. Both at 300 V/m, and at the maximum strength of 436 V/m, the speed is significantly higher than under all other conditions (*p <* 0.001; asterisks, ANOVA on ranks, all-pairwise comparisons using Dunn’s test).

**Fig 2.**
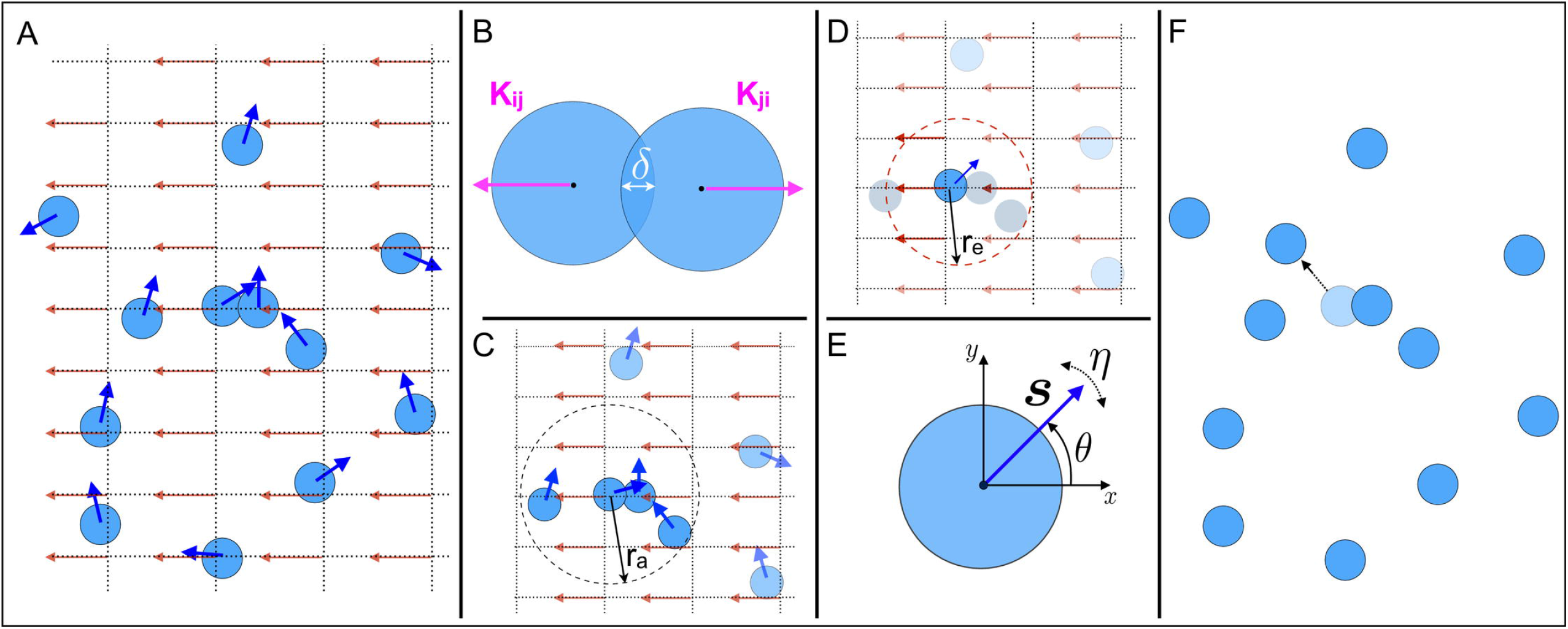
Theoretical model for cell migration in a DC electric field. (A) Osteoblasts in the cell culture chamber exposed to the DC electric field, are modelled as active particles (light blue coloured disks) of radius *R*, and, their motion at any time *t* is described by the velocity 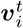(*i* is the index of the cell). The dark blue coloured arrows laid over the circular disks are the unit vectors 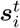 of the cell velocity with the magnitude *v*_0_ at time *t*. The model takes into account both cell-cell and cell-electric field interactions. These interactions can influence the cell velocity. Cell-cell interactions involve finite-volume exclusion and migration orientation alignment. (B) When two cells overlap, cell *i* experiences a displacing force *K_ij_* from each of its neighbouring cell *j*, where *i* and *j* are the cell indices. The magnitude of such a force is proportional to the degree of overlap *δ*. (C) The migration direction of a cell can be influenced by its neighbouring cells located within the radius *r_a_*, taken from the cell’s center. Such an interaction re-orients the migration direction of a cell to the average direction of migration of neighbouring cells. (D) Each individual cell also experiences a force due to the electric field. Electric field is defined on a discrete two-dimensional grid underlying the space in which the cells move. The cell experiences the average force from the electric field at all the grid points that lie within the radius *r_e_*, taken from the cell’s center. (E) The limited precision in cellular sensing of directional alignment is captured by an angular white noise term whose strength is given by *η*. (F) The resultant direction *θ* of cells migration is the cumulative effect of the cellular interactions described in B-E. The model at each time step calculates and updates the position of each cell (shown by dotted arrow for the cell under consideration) for the next time step.

Comparing cell migration velocities (plotted as sectors of polar plots) without stimulation (Fig 1 D), to those with weak (160 V/m; Fig 1 E) or strong stimulation (436 V/m; Fig 1 F), one can appreciate that the directionality of migration shifts from random, covering all sectors of the plot (Fig 1 D) to exclusively anodal, covering only the anodal sectors (Fig 1 F), with increasing field strength. At the same time, also the speed of the cells appears to shift from lower speeds with a maximum of 8-10 *μ*m/h (green hues in Fig 1 D corresponding to ~ 2.5 % of the cells) to a maximum of 16-18 *μ*m/h (red sectors in Fig 1 F corresponding to ~ 12 % of the cells).

To address the question of a possible correlation of speed and field strength, we quantified the cell migration speed of all cells in all experiments under different stimulation strengths. As shown in Fig 1 G, migration speed under DC-stimulation is significantly different from sham stimulation conditions only at 300 V/m and 436 V/m (p¡ 0.001, ANOVA on ranks with Dunn’s all-pairwise comparisons). Considering all values and calculating a linear regression (blue line in Fig 1 G), there is thus a weak correlation between field strength and migration speed, with a regression coefficient of *R* = 0.516.

### Modeling electrotactic collective osteoblast cell migration

We describe the *in vitro* motility behaviour of individual cells that are subject to external DC electric field. The main components of our model are (i) the ability of the cells to interact with the other cells, and, (ii) the ability of the cells to interact with the external electric field. Cell-cell interaction involves two types of forces: short-range repulsive forces, and the alignment of the direction of motion with the cells’ local neighbours. The force at the short distances, through soft-core repulsion, ensures that cells do not overlap. We also include in our model the influence of each cell’s local neighbours on the direction of its migration. Such cell-to-cell interactions are certainly playing a role in high-density tissue culture via cell-cell contacts. However, since mechanical or signalling cues are at least conceivable also in 2D cell cultures without direct cell contacts, we introduce this factor in the model to study the possible role of such interactions in our experiments. Finally, we also consider the interaction of cells with the applied DC electric field.

### The cell

Each cell is modeled as a circular disk of radius *R* which can migrate in two spatial dimensions with an active speed of *v*_0_. The state of each cell *i* is characterised at time *t* by its position 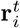, described through the coordinates 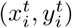, and its migration velocity 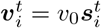, where, *v*_0_ is the cell migration speed and 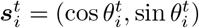 is the unit vector representing the direction of migration, with *θ* being the angle that the cell makes with the horizontal axis of the laboratory frame. The direction *θ* that each cell takes at any consecutive time depends not only on its direction of motion in the immediately preceding time, but also on the forces acting on the cell. The total force acting on the cell *i* results from cell-cell interactions and cell interaction with the applied DC electric field. These forces are discussed in more detail in the following sections.

### Cell-cell interactions

We consider two types of cell-cell interactions in our model. The cell-cell interaction due to finite-volume exclusion and the cell-cell interaction resulting from cell orientational alignment with its neighbours. Each cell is assumed to occupy a finite area in the cell culture medium in which it is placed. To avoid cell overlaps, we include, at each time *t*, repulsive force 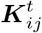 that is proportional to the degree of overlap between two cells and is given by,

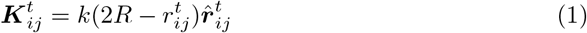

with,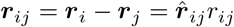 and *k* a force constant. *r_ij_* is the euclidean distance between two cells *i* and *j* and is calculated as 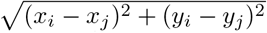. The total repulsive force acting on cell *i* at time *t*, denoted by 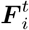 is,

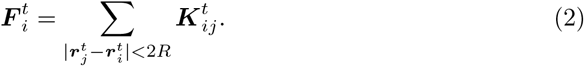

The directional alignment of cells with its proximal neighbours is given by,

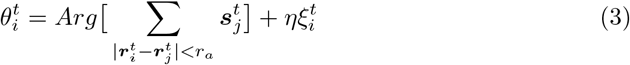

and is only hampered by an angular white noise uniformly distributed in [−*π, π*] with 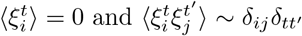 and whose strength is given by *η*. The first term in Eq (3) returns the average of the angle defining the orientation of the vector 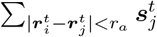, where the sum extends only to those cells which are within the interaction radius *r_a_* of cell *i*.

### Cell-electric field interaction

The electric field is defined on a regular square lattice underlying the domain in which the cells are migrating. Each grid point is specified by coordinates (*p, q*). The electric field at the grid point located at (*p_k_, q_k_*) is characterised by the vector ***E**_k_* = *E_0_ **e_k_***, where, ***e**_k_* = (cos Θ_*k*_, sin Θ_*k*_) is the unit vector representing the polarity of the electric field, and, *E*_0_ corresponds to the electric field magnitude of electrical stimulations in the experiments. Θ_*k*_ is the angle that the electric field vector at (*p_k_, q_k_*) makes with the horizontal axis. Cell *i* experiences an effective electric field which is the average of the electric field on all the grid points that lie in the region within the radius *r_e_* from the center of cell *i*. The net electrical force acting on cell *i* is given by,

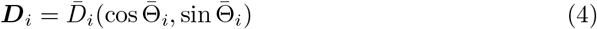

where, 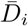 and 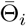 are the mean strength and the mean orientation of the force due to the electric field acting on the cell at the location (*x_i_, y_i_*).

The position of each cell *i* is updated in the next time step, after calculating the resultant angle *θ* due to all the interactions, including the cell neighbour orientation alignment, by the following scheme:

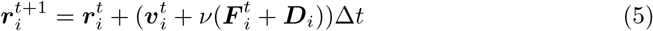

where, *ν* is a friction factor that is associated with the cell substrate interaction, and, ∆*t* is the time step.

### Simulation details

We simulate the motility behaviour of *N* = 35 cells, as in our experiments there are approximately 30-40 cells in a single field of view. Cells are initially randomly distributed in a circular region within the spatial domain representing the stimulation chamber. Osteoblast cells are roughly 100 *μm* in diameter, considering all cells extensions, and, we use this to define the cell radius *R*, which is one length unit in our simulations. The cell radius *R* is assumed to be the basic length scale in these simulations. Time steps are seperated by ∆*t* which is set to 1. The parameters of the model and their values used in these simulations are listed in Table 1. The active speed of cells is 0.1*R* per time step. The time parameters in the simulations are scaled such that the speed of the unstimulated cells corresponds to the average cell speed in the experimental sham case, which is ~ 2 - 3*μ*m*/* h. The magnitude of the electric field *E*_0_ = 0.014 corresponds to 160 V/m in experiments, and is chosen such that the cell speed, in this case, is approximately equal to the experimental value, i.e ~ 4*μm/h*. The magnitude of the electric field corresponding to 436 V/m in experiments is obtained by simply multiplying 0.014 by the factor 2.75, as 436*/*160 = 2.75. At the start of each simulation, we specify the initial positions 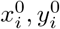, initial speed 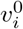 and the orientation 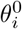 of each cell *i*. The initial orientation is distributed randomly in the range [0,2*π*]. At each time step for each cell we identify cells which are less than a distance of 2*R* apart. From this we calculate the force due to volume exclusion acting on each cell from its neighbouring cells, as given by Eq (1) and Eq (2). We also determine all the grid points of the underlying grid, on which the electric field is defined, that are within the radius of *r_e_* of each cell and calculate the mean electric field. This constitutes the net force due to the electric field ***D**_i_* acting on each cell *i*, as given by Eq (4). Experiments show that the cell migration is anode-directed. We incorporate this into our model by assigning a polarity to the mean electrical force experienced by the cell, and it is opposite to the applied electric field, i.e 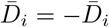. In addition, we also determine for each cell all its neighbouring cells that are located within the radius *r_a_*, and calculate the mean orientation of all those cells. Each cells’ orientation is updated by its mean orientation, to which a weak noise *η* = 0.05 is added, as given by Eq (3). Finally, the positions of each individual cell is then updated using the Eq (5).

**Table 1.**
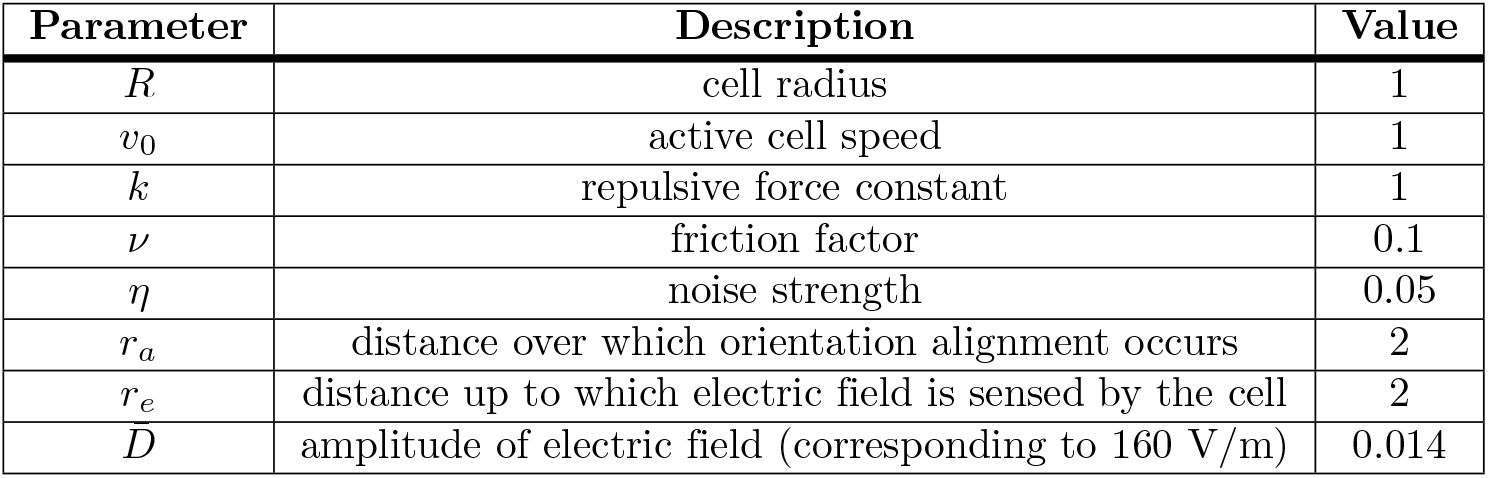
List of all the model parameters, their notation, description and value (dimensionless).

### Migratory behaviour of osteoblasts in DC electrical field

To study the influence of externally applied DC electrical field on the migratory behaviour of human osteoblast we simulated *N* = 35 migrating cells with and without DC electrical stimulation for 130 time steps. We have verified, through multiple runs of the simulations, that our results are qualitatively invariant of random initial conditions and stochastic angular fluctuations in the simulations, S1 Fig. We use periodic boundary conditions, to reflect the experimental conditions in which cells are placed in the center, and thus far from the boundaries, of the stimulation chamber. Fig 3 left column shows the positions of all cells at the final time step, in the case of no electrical stimulation, Fig 3 A, and in the case of DC electrical stimulation with field amplitudes of 0.014 and 0.038, Fig 3 B and C, respectively. Electric field amplitude of magnitude 0.038 in simulations corresponds to the maximum field strength of electrical stimulation in experiments, i.e., 436 V/m, Fig 1 G. Fig 3 D-F (middle column) presents the individual cell trajectories at each time step in the case of no electrical stimulation, Fig 3 D, and in the case of electrical stimulation with different field amplitudes of 0.014 and 0.038, Fig 3 E and F, respectively. The velocity of cell migration, calculated from the initial and the final time step, is shown in Fig 3 G-I (right column) as polar plots for the case without electrical stimulation, Fig 3 G, and with electrical stimulation of different field amplitudes, i.e.,0.014 and 0.038, Fig 3 H and I, respectively. Each polar plot shown in Fig 3 G-I, is the cumulate of 10 separate runs of the simulation. Initial velocity of each cell and the noise in cell velocity at each time step are random, this renders robustness to the polar plot distributions.

**Fig 3.**
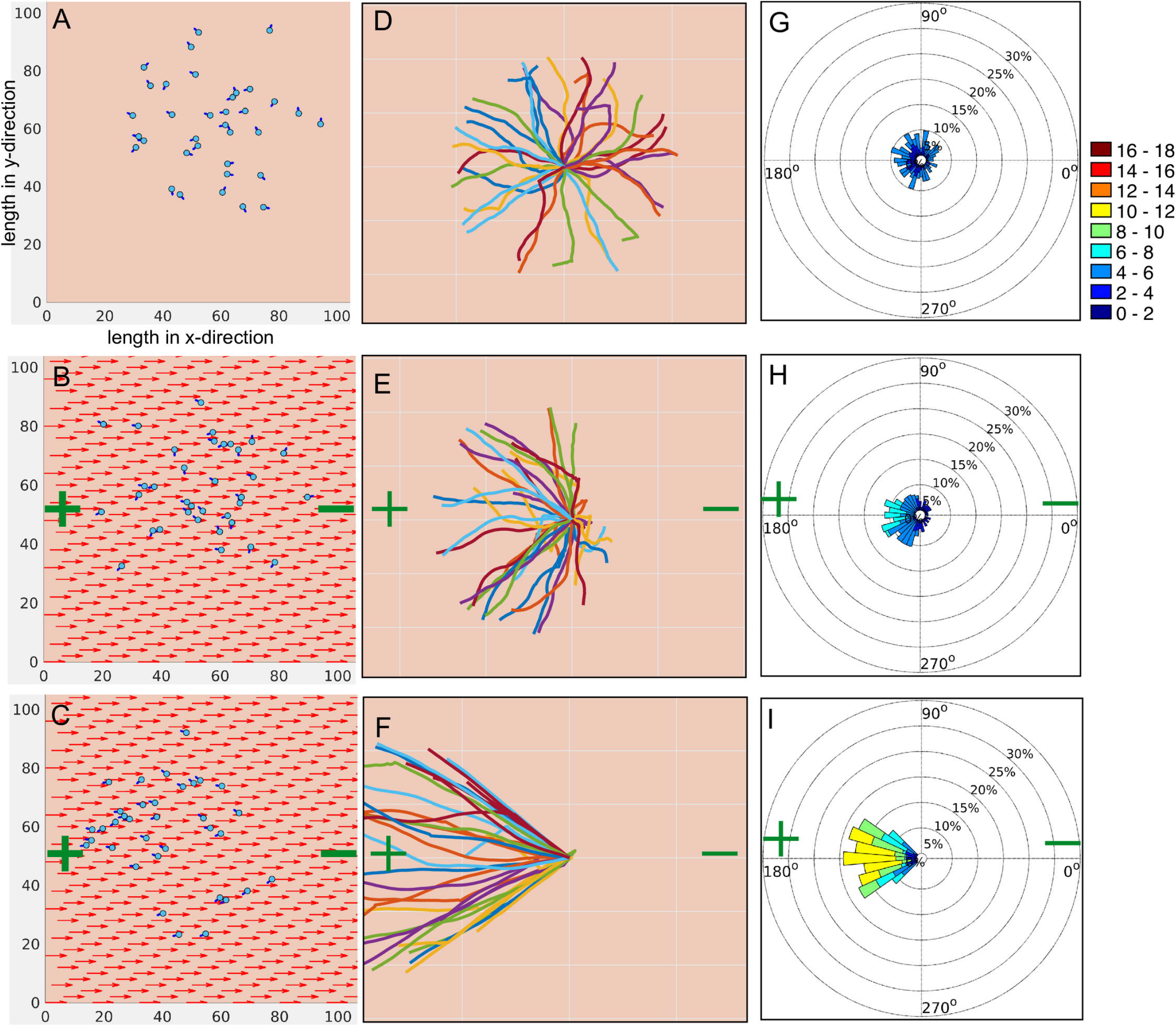
Simulation of cell migration model in DC electrical field. Each simulation consists of 35 cells, initially randomly distributed in a circular region around the center of the domain of size 120×120. Osteoblasts are modelled as light blue coloured circular disks of radius R=1 with randomly oriented initial migration velocity represented by dark blue arrows. The polarity of the DC electric field is denoted by green coloured plus and minus symbols, and, the electric field vector is represented by red arrows. The model is simulated for 130 time steps. (A-C) Final positions of individual cells in the case of, A unstimulated cells, and, DC electrical stimulation of strength 0.014 and 0.038, shown in B and C, respectively. Electric field strength of 0.038 in simulations corresponds to the maximum electric field strength of 436 V/m in experiments. (D-F) Trajectories of individual cells corresponding to the three cases shown in A-C, respectively. Cell positions are adjusted such that all the trajectories originate from *x* = 0 and *y* = 0 at *t* = 0. (G-I) Polar plots showing the velocity of cell migration, taking into account only the initial and the final time step. G shows the polar plot for the unstimulated case, whereas H and I present polar plots for 160 V/m and 436 V/m, respectively. Each polar plot is a cumulate of data from 10 separate simulation runs, where each simulation consists of 35 cells.

In the absence of electrical stimulation the cells move, as expected, in all directions, Fig 3A. Trajectories of individual cells show that, over time, all cells collectively explore the space homogeneously, Fig 3B, a feature which is also reflected in the polar plots, which are constructed, similar to the experiments, based only on the initial and final time steps Fig 3G. The mean cell speed in this case is ~ 3*μ*m/h. However, when DC electrical field of amplitude 0.014, which corresponds to 160 V/m, is applied, cells start exhibiting a directional migration towards the anode 3B. Individual cell trajectory plot shows that although the final position of the majority of the cells is towards the anode, few cells still migrate towards the cathode, albeit much shorter distances than the anodally migrated cells, Fig 3E. The polar plot, showing velocity of cell migration, clearly presents the modulation of the orientation of migration by external field, Fig 3H.

Following the trajectories of individual cells also reveals that cell migration is not instantaneously switched in the direction of anode. Cells respond to the applied electrical field by gradually changing their directionality of migration. Initially most of the cells move in a random manner, however, at later times, gradually turn towards the anode. This delayed response in eventual anode directed motility of cells is because the electrotactic migration speed *μ**D*** is much weaker than the active cell migration speed ****v****. Cells display a highly directional motion towards the anode with increasing strength of the electric field *E*_0_, Fig 3C. Cell migration in this case shows a much faster re-orientation and much persistent motion towards anode 3F. Fig 3I shows that not only the direction of the motion is influenced by increasing field strength, but also the velocity of the cell migration. The maximum cell speed in this case even reaches up to 10-12 *μ*m/h, Fig 3I. To better quantify the changes in the collective cell migratory behaviour we calculate, in both experiments and simulations, the directionality order parameter Φ, which reflects how well cell movements have aligned with the electric field and directed towards the anode, and is given by,

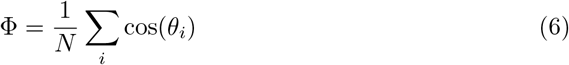

where, *N* is the total number of cells and the sum is over the cosine of migration direction of individual cells *θ_i_*. Φ can vary between 1 (towards cathode) and −1 (towards anode) and Φ ≃ 0 corresponds to random cell movement. Our results show that for the listed choice of parameters, the directionality order parameter Φ obtained from the model simulation matches very closely with the experiments, Fig 4A.

**Fig 4.**
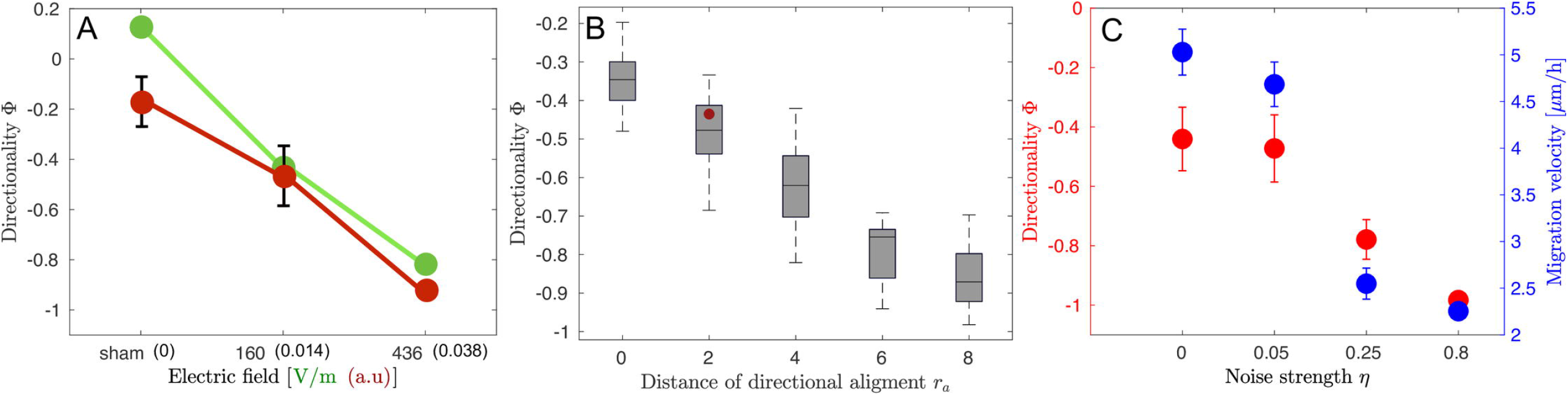
Influence of the electric field strength, radius of alignment and noise on the directionality of cell migration. (A) Comparison of the directionality order parameter Φ obtained from the simulations (red colour filled circles connected by red lines) with experiments (green colour filled circles connected by green lines) corresponding to the three different cases, i.e no stimulation, stimulation with field strength 160 V/m and 436 V/m, respectively. The electric field strength in simulations is shown in arbitrary units (a.u), where 0.038 (in brackets) corresponds to the maximum field strength of 436 V/m in experiments. Each value of directionality obtained from simulations is the average of 10 separate simulation runs. Each simulation consists of 35 cells. Error bars in simulation data show the standard deviation in Φ. (B,C) Parameter sweep was performed to study the influence of the radius of directional alignment and noise strength on the directionality of migration in the case of electrical stimulation of amplitude 0.014, which corresponds to 160 V/m in experiments. B shows the box and whisker plot of medians (horizontal lines) of directionality of migration Φ vs. distance of directional alignment *r_a_*. The values of all other parameters, except *r_a_*, are as mentioned in Table 1. Cells show higher directedness Φ in their migration towards anode with increasing distance *r_a_* over which directional alignment occurs. The red dot is the experimental value for the directionality in the case of electrical stimulation of strength 160 V/m. Whiskers denote 25-75 percentiles of data distribution. Φ = −1 corresponds to fully directed movement towards the anode, which is located at 180° in Fig 1 D-F and Fig 3 G-I. C presents directionality order parameter Φ (red) and migration velocity (blue) vs. noise strength *η*. The values of all other parameters, except *η*, are as mentioned in Table 1. Increasing noise strength leads to higher directedness in cell movement towards the anode Φ ~ −1. However, for the same values of noise strength, migration velocity decreases with increasing noise strength. Error bars show the standard deviation in the directionality and the migration velocity for different values of noise strength obtained from simulations.

Our results shown in Fig 3, and, Fig 4A, reproduce the following experimental observations (i) in the absence of electrical stimulation, which corresponds to the experimental sham case, directional migration of cells is not observed, i.e, cells do not collectively move in a preferred direction, (ii) alignment of the directionality of cell migration depends on the strength of the applied electrical field, (iii) average cell velocity increases with the field strength.

### Influence of radius of alignment and noise on cell migration

In our model the collective behaviour results from the directional alignment of individual cells with each other. This is controlled by the model parameter *r_a_*, which is the distance over which the cell aligns its direction of migration with its neighbours, and *η*, which is the strength of the fluctuation in the direction of migration of individual cell. In the simulation results discussed in the preceding section, Fig 3, we considered *r_a_* = 2*R*, i.e., the orientation alignment occurs only when cells touch each other. In order to understand how the two model parameters *r_a_* and *η* affect the migratory behaviour of electrically stimulated osteoblast cells, we perform a parameter sweep study with fixed electrical stimulation of strength 0.014 (which corresponds to 160 V/m in experiments). To study the influence *r_a_* and *η* on collective cell migration, we obtain directionality Φ for different values of *r_a_* and *η*, as shown in Fig 4A and B, respectively. The values of all the other parameters are as mentioned in Table 1. Our results show that, even in the case of weak electrical stimulation, which corresponds to 160 V/m in experiments, with increasing *r_a_* the cells move in a more directed manner towards the anode, i.e, Φ approaches the value of −1, as shown in Fig 4A. Cell movement also displays a higher directedness with increasing noise strength *η*, which is unexpected, Fig 4B and S2 Fig. Although cells move in a more directed manner towards anode, the cell migration velocity, on the contrary, decreases with increasing noise strength, Fig 4B. Taken together these results suggest that the parameters, i.e, *r_a_* and *η* can significantly alter the dynamics of cell migration and give rise to collective electrotactic motion of osteoblast cells.

## Discussion

The migration of osteoblasts, which plays a key role in bone regeneration, can be modulated by external electrical stimulation [30]. This offers an attractive approach towards building electrically active implants for effective tissue regeneration [31–33]. In the present paper, we presented a computational model to study (i) the migratory behaviour of osteoblasts, and, (ii) the consequences of the application of external electrical field on their migration. The model was used to study the collective behaviour of many cells in *in vitro* experiments where primary human osteoblasts placed in electrotaxis chamber were stimulated by DC electric field. For this purpose, were-analysed the galvanotactic migration of human osteoblasts exposed to DC-electric field stimulation at different field strengths from a previous study published in [14], now using single-cell rather than clustered data. As observed in our previous paper [14], we confirmed that field exposition leads to migratory directionality towards the anode, and elucidate that the migratory speed distribution ranges from 2-18 *μ*m/h, with significantly higher speeds of migration than unstimulated cells at DC-field strengths of 300 and 436 V/m. Using this single-cell analysis approach, beyond our initial findings in the cited paper using pooled data (i.e. stimulated vs. unstimulated only), we show that the directionality thus actually significantly depends on the field strength, with random migration without stimulation, ~ 65% anodal migration at low (160 V/m) and exclusively anodal migration at highest field strength (436 V/m). Our detailed cell-by-cell analysis also shows that, although directionality of cell migration clearly correlates with the strength of the applied electric field, there is only a weak correlation of migratory speed and electric field strength, a correlation which could not be seen in the pooled analysis of our previous paper.

To explain these experimental observations, we modeled each cell as an active agent whose movement is influenced by its own interactions with other cells, external electric field and stochastic switching in the direction of migration. The model takes into account the force experienced by the cell due to the applied DC electric field. We also considered two types of inter-cellular interactions: in addition to the nearest neighbour interaction that ensures finite-volume exclusion by penalizing cell overlaps, cells also interact with other cells via a velocity alignment mechanism. Although specific molecular mechanisms underlying these interactions remain unclear, two important questions can be addressed by the current simulation study: (i) Does directionality also depend on interaction among neighbouring migrating cells, and if so, how large is this interaction radius, (ii) Do directionality and migration speed depend on the accuracy of the putative cellular field sensing mechanism, i.e. in which way does a noise factor influence migration directionality and migration speed?

Our results show that the motility behaviour of cells is influenced by the distance over which the cell aligns with its neighbours, stochastic switching in the direction of migration and the strength of applied electric field. The simulations in the present paper closely match the experimentally observed weak correlation between migration speed and the applied electric field, and are more realistic than previously published ones [17], which predicted speed ranges from 1.8 to 4.0 *μ*m/s, i.e. nearly tenfold the maximum observed by us. As discussed previously, migration at such high speeds probably finds its limitations in adhesive forces acting on the cells on the one hand, and rate-limiting factors such as actin conformational change being limited by temperature and Ca^2+^ dynamics [34, 35]. We performed a quantitative comparison of the directionality order parameter obtained from simulations with experiments as shown in Fig 3(J), where directionality angles Φ of experimental values and simulations practically overlap. As the simulation results show, varying *r_a_* from 0 (i.e the case with no inter-cellular interactions) to 8 (i.e. the case with inter-cellular interactions between two cells extending to distances of four cell diameters), the directionality of ~ −0.45 for electrical stimulation of strength 160V/m, as found in our experiments, best matches with a value of *r_a_* of 2. These results suggest that the interactions between cells only in direct contact likely lead to parallel anodal movement. The mechanism of this interaction could be speculated to rely on e.g. osteoblast binding via cadherin, an interaction known to be important for morphogenesis of osteoblasts, and subsequent modulation of actin function [36, 37]. Long-distance effects, mediated by e.g. molecules secreted from the cells, tension changes within the collagen coating, or distortion of the electric field by the neighbouring cell are, in turn, unlikely to be important for osteoblasts.

Our results also show that stochastic orientational switching can significantly alter cellular electrotactic motility behaviour. In this case, a perfectly directed motion towards the anode is achieved for very high fluctuation strengths, which appears to be counter-intuitive since one would expect that for higher angular fluctuations the accuracy of directional movement aligned with the electric field decreases. Varying *η* in our simulations from 0 to 0.8, the directionality of ~ −0.45 in our experiments is in line only with a very low degree of noise (around 0.05, which corresponds to fluctuations of ~ 10° in the direction of cell migration), but not commensurate with values of *>* 0.25. The experimental migration speed found to be in the range of 2 to 12 *μ*m/h would also cover the simulated value of ~ 4.75 *μ*m/h at *η* = 0.05. It is, however, conceivable, that other cell types do show more influence of noise (arguably reflecting e.g. less mechanical interactions with the substrate, varying cell shape influences, or different field sensing or signalling mechanisms). What remains to be explained is the seemingly paradoxical result that higher fluctuation levels should lead to higher accuracy in directionality. Our hypothesis would be that higher fluctuation actually raises the probability of cell-to-cell interactions, which in turn will lead to common field alignment. If this hypothesis holds true, such movement would lead to field orientation of cells with higher accuracy but lower speed due to frequent corrective movements. Although experiments clearly are needed to validate this hypothesis, it is interesting to note that at the highest stimulation strength of 436 V/m, those cells which are best aligned to the field and directed towards anode do not belong to the fastest subset of cells (which are, indeed, 10°-30°off the “ideal” orientation; see Fig 1F.

## Conclusion

Our computational model provides a framework for studying cell migration under the influence of the DC electric field, and, elucidating the rules and the role of individual cell interactions, with other cells and with their physical environment. This model is also relevant to study the influence of additional factors on cell migration, such as the cell density and other modes of electrical stimulation, e.g. alternating current stimulation. The model we present here allows for easy integration of additional details, as more data becomes available. Our approach could serve as a tool to not only test existing hypotheses of electrotactic cell migration but also predict migratory behaviour under perturbation conditions, and thus bridge the gap between single cell and collective response in more effective manner.

## Materials and methods

In this study, data on cell migration of human osteoblasts under DC-electrical stimulation were re-analysed using a previous set of experiments [14]. Cell cultivation and stimulation methods are detailed in this paper, and given in brief below:

### Cell culture

Human osteoblasts were isolated from femoral heads of patients (*n* = 14) undergoing a total hip replacement. Patients gave consent and the study was approved by the local ethics committee (permit A 2010-10). Osteoblasts were isolated from cancellous bone as previously described [38]. Isolated cells were cultured in Dulbecco’s Modified Eagle Medium (Pan Biotech, Aidenbach, Germany) supplemented with 10% fetal calf serum, 1% amphotericin B, 1% penicillin-strepto-mycin and 1% hepes- buffer under standard cell culture conditions (5% CO_2_ and 37°C). Ascorbic acid (50 *μ*g/ml), *β*-glycerophosphate (10 mM), and dexamethasone (100 nM) (Sigma Aldrich, St. Louis, MO, US) were added to cell culture medium to maintain osteoblast phenotype. For cell migration experiments cells in passage three were used.

### DC electrical stimulation chamber and experimental procedure

To study migration of osteoblasts in electric fields, we used a two-part stimulation chamber described in [14]. Before each use, both chamber parts were cleaned with 70% ethanol, washed with a mild detergent and rinsed extensively with distilled water before steam sterilization. Coverslips (24 × 50 mm) for seeding osteoblast cultures were coated with rat tail collagen (Advanced Biomatrix, San Diego, CA, USA) by incubation of 50 *μ*m/ml rat tail collagen diluted in sterile 0.1% acetic acid for 1 h. Coverslips were positioned in a groove in the upper chamber part and edges sealed with silicon paste (Korasilone, Obermeier GmbH, Bad Berleburg, Germany). Upper and lower chamber parts were bolted by 12 screws to ensure tight contact and prevent leakage and chambers were exposed to UV light for sterilization. After this sterilisation treatment, remaining solution was aspired and coverslips were washed twice with phosphate buffered saline (Biochrom, Berlin, Germany) before cell seeding. A total of 2 × 10^3^ osteoblasts were seeded per chamber and cells were allowed to adhere for 30 min. Afterwards, coverslips were washed twice with medium to remove non-adherent cells. Chambers were then sealed with a top coverglass, and silicon paste and cells accommodated to chamber overnight. For DC-stimulation, silver/silver chloride electrodes were placed into outer reservoirs separated from cell area to avoid electrochemical reactions within the tissue chamber. Current was conducted to the cell chamber using agar bridges (silicon tubes, length 120 mm, inner diameter 5 mm) consisting of 2% agarose (TopVision agarose, ThermoScientific, Waltham, MA, US) in Ringer’s solution (Braun, Melsungen, Germany). Current was applied to electrodes for 7 h via crocodile clamps using a DC power supply (Standard Power Pack P25, Biometra, Göttingen, Germany). To maintain constant stimulation, voltage was measured directly at the borders of the cell area (electrode distance 24 mm) using a multimeter (Voltcraft VC220, Conrad Electronic, Wollerau, Switzerland) and adjusted during the experiments. Each of the experiments was conducted with one cell culture being divided to obtain sham stimulation group as control, and a DC-stimulation group for the respective field strength used. Electric field strengths were 160, 300, 360, 426 and 436 V/m.

### Migration analysis

For the analysis, all cells from the sham groups were pooled as one control. Thus, a total of n=177 (sham), 34 (160 V/m), 35 (300 V/m), 26 (360 V/m), 43 (426 V/m) and 33 (436 V/m) cells were analysed. For this, photographs were taken at 8 fields of view evenly distributed over the cell area at beginning (Fig 1A) and end time points (Fig 1B) with a Leica DMI 6000 and LAS X software during the 7-hour stimulation, or sham stimulation, procedure. The pairs of photographs were then aligned manually, and merged, taking external markers as reference points (Fig 1C). To quantify migration within the electric field, segmentation of the cell shape, including cell extensions, was performed manually using Image J software (NIH) for each cell that could be identified in both the time points (see yellow coastlines in Fig 1C), i.e 0 h and 7 h after DC stimulation. The segmentation procedure yielded, among other features, the coordinates of the cell centroid. Using the coordinates of the cell centroid at these two time points, i.e., 0 h and 7 h, the distance and orientation of migration was calculated for each cell. The migration distance was defined as 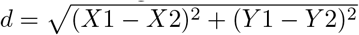, where *X*1, *Y* 1 and *X*2, *Y* 2 represent the coordinates of the cell centroid at 0 h and 7 h after DC stimulation, respectively Fig 1C. The migration angle was defined as 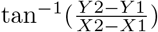. Using the migration distance and orientation we obtained a migration plot for each case, which could be depicted in a polar coordinate system, as shown in Fig 1D-F. The anode in the polar plots of DC stimulated experiments is located at 180°angle, Fig 1E and F. For better comparison of all experiments, we binned the migration angles into 36 sectors of 10° each, and classified migration speeds in a scoring system. Thus, the migration angle was calculated starting from the original cell position, and angles were assigned to the 36 sectors, where sector 10-18 (90-180°) and 18-26 (180-270°) represent anode-directed migration, while sectors 1-9 and 27-36 (0-90 and 270 to 360 °) represent cathode-directed cell migration. To construct polar plots (Fig 1 D-E) illustrating both migration direction and velocity, the migration speed of single cells was colour coded from 0 to 18 *μ*m/h in 9 groups of 3 *μ*m/h bins. The relative sector lengths denote the percentage of cells migrating at a certain speed range.

## Acknowledgments

This research was funded by Deutsche Forschungsgemeinschaft (DFG, German Research Foundation), under the grant SFB 1270/1-299150580. We thank Ms. Doris Hansmann for providing the primary human osteoblasts. We thank Kai Budde, for his helpful and constructive comments on our manuscript.

## Supporting information

**S1 Fig.**
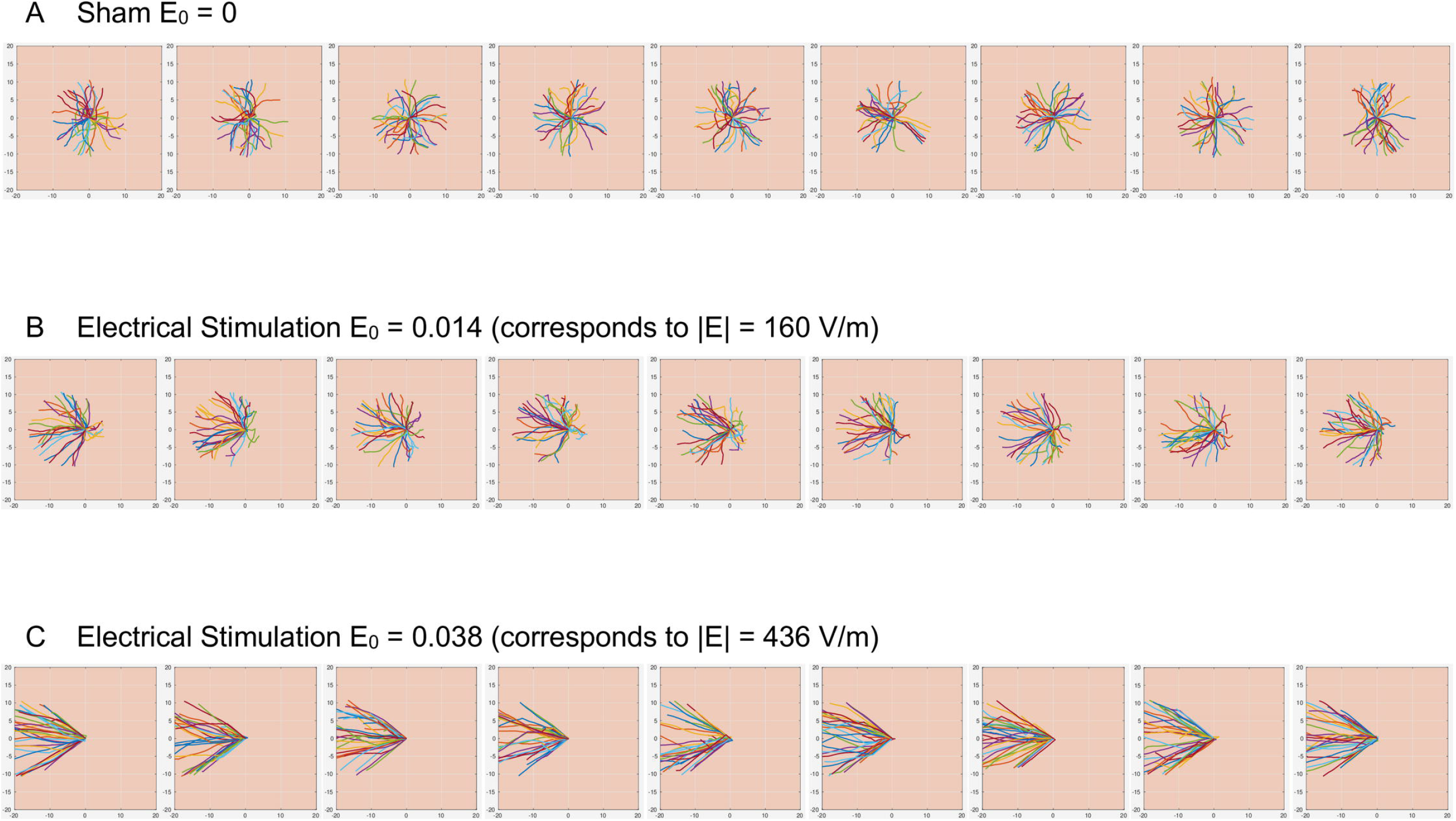
Individual cell trajectories for multiple simulation runs. Results of ten separate simulation runs for the case: (A) no electrical stimulation, which corresponds to experimental sham, (B) stimulation with electrical field amplitude of 0.014, which corresponds to the experimental field stimulation strength of 160 V/m, and, (C) stimulation with electrical field amplitude of 0.038, which corresponds to the experimental field stimulation strength of 436 V/m.

**S2 Fig.**
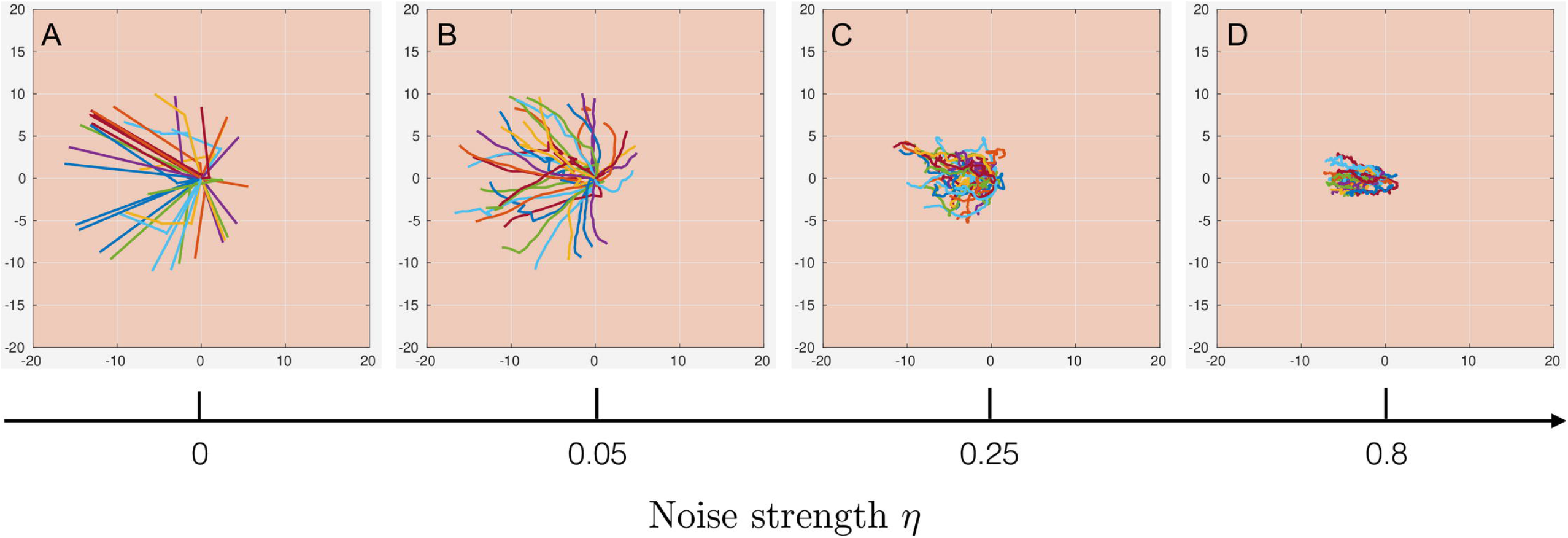
Influence of noise on cell migration pattern. The trajectories of individual cells in the simulation case corresponding to the experimental electrical stimulation of 160 V/m with increasing noise strength in the direction of migration, (A) 0, (B) 0.05, (C) 0.25, (D) 0.8.

## Notes

### Competing Interest Statement

The authors have declared no competing interest.

## References

1. Simpson MJ, Lo KY, Sun YS. Quantifying the roles of random motility and directed motility using advection-diffusion theory for a 3T3 fibroblast cell migration assay stimulated with an electric field. BMC Systems Biology. 2017;doi:10.1186/s12918-017-0413-5.

2. Lara Rodriguez L, Schneider IC. Directed cell migration in multi-cue environments; 2013.

3. Wu D, Lin F. A receptor-electromigration-based model for cellular electrotactic sensing and migration. Biochemical and Biophysical Research Communications. 2011;doi:10.1016/j.bbrc.2011.07.004.

4. Zajdel TJ, Shim G, Wang L, Rossello-Martinez A, Cohen DJ. SCHEEPDOG: Programming Electric Cues to Dynamically Herd Large-Scale Cell Migration. Cell Systems. 2020;doi:10.1016/j.cels.2020.05.009.

5. Liang Y, Tian H, Liu J, Lv YL, Wang Y, Zhang JP, et al. Application of stable continuous external electric field promotes wound healing in pig wound model. Bioelectrochemistry. 2020;doi:10.1016/j.bioelechem.2020.107578.

6. Cho Y, Son M, Jeong H, Shin JH. Electric field-induced migration and intercellular stress alignment in a collective epithelial monolayer. Molecular Biology of the Cell. 2018;doi:10.1091/mbc.E18-01-0077.

7. Lin BJ, Tsao SH, Chen A, Hu SK, Chao L, Chao PHG. Lipid rafts sense and direct electric field-induced migration. Proceedings of the National Academy of Sciences of the United States of America. 2017;doi:10.1073/pnas.1702526114.

8. Saltukoglu D, Grünewald J, Strohmeyer N, Bensch R, Ulbrich MH, Ronneberger O, et al. Spontaneous and electric feld-controlled front-rear polarization of human keratinocytes. Molecular Biology of the Cell. 2015;doi:10.1091/mbc.E14-12-1580.

9. Hart FX, Palisano JR. Glycocalyx bending by an electric field increases cell motility. Bioelectromagnetics. 2017;doi:10.1002/bem.22060.

10. Zhao M, Pu J, Forrester JV, McCaig CD. Membrane lipids, EGF receptors, and intracellular signals colocalize and are polarized in epithelial cells moving directionally in a physiological electric field. The FASEB journal: official publication of the Federation of American Societies for Experimental Biology. 2002;doi:10.1096/fj.01-0811fje.

11. Zhao Z, Watt C, Karystinou A, Roelofs AJ, McCaig CD, Gibson IR, et al. Directed migration of human bone marrow mesenchymal stem cells in a physiological direct current electric field. European Cells and Materials. 2011;doi:10.22203/eCM.v022a26.

12. Sato MJ, Kuwayama H, Van Egmond WN, Takayama ALK, Takagi H, Van Haastert PJM, et al. Switching direction in electric-signal-induced cell migration by cyclic guanosine monophosphate and phosphatidylinositol signaling. Proceedings of the National Academy of Sciences of the United States of America. 2009;doi:10.1073/pnas.0809974106.

13. Guido I, Diehl D, Olszok NA, Bodenschatz E. Cellular velocity, electrical persistence and sensing in developed and vegetative cells during electrotaxis. PLoS ONE. 2020;doi:10.1371/journal.pone.0239379.

14. Rohde M, Ziebart J, Kirschstein T, Sellmann T, Porath K, Kühl F, et al. Human Osteoblast Migration in DC Electrical Fields Depends on Store Operated Ca2+-Release and Is Correlated to Upregulation of Stretch-Activated TRPM7 Channels. Frontiers in Bioengineering and Biotechnology. 2019;doi:10.3389/fbioe.2019.00422.

15. Gruler H, Nuccitelli R. The Galvanotaxis response mechanism of keratinocytes can be modeled as a proportional controller. Cell Biochemistry and Biophysics. 2000;doi:10.1385/cbb:33:1:33.

16. Schienbein M, Gruler H. Langevin equation, Fokker-Planck equation and cell migration. Bulletin of Mathematical Biology. 1993;doi:10.1007/BF02460652.

17. Vanegas-Acosta JC, Garzón-Alvarado DA, Zwamborn APM. Mathematical model of electrotaxis in osteoblastic cells. Bioelectrochemistry. 2012;doi:10.1016/j.bioelechem.2012.08.002.

18. Waters CM, Bassler BL. Quorum sensing: Cell-to-cell communication in bacteria; 2005.

19. Thurley K, Wu LF, Altschuler SJ. Modeling Cell-to-Cell Communication Networks Using Response-Time Distributions. Cell Systems. 2018;doi:10.1016/j.cels.2018.01.016.

20. Barton DL, Henkes S, Weijer CJ, Sknepnek R. Active Vertex Model for cell-resolution description of epithelial tissue mechanics. PLoS Computational Biology. 2017;doi:10.1371/journal.pcbi.1005569.

21. Henkes S, Kostanjevec K, Collinson JM, Sknepnek R, Bertin E. Dense active matter model of motion patterns in confluent cell monolayers. Nature Communications. 2020;doi:10.1038/s41467-020-15164-5.

22. Bittig AT, Jeschke M, Uhrmacher AM. Towards modelling and simulation of crowded environments in cell biology. In: AIP Conference Proceedings; 2010.

23. Vicsek T, Czirk A, Ben-Jacob E, Cohen I, Shochet O. Novel type of phase transition in a system of self-driven particles. Physical Review Letters. 1995;doi:10.1103/PhysRevLett.75.1226.

24. Bhattacharya K, Vicsek T. Collective decision making in cohesive flocks. New Journal of Physics. 2010;doi:10.1088/1367-2630/12/9/093019.

25. Szabó B, Szöllösi GJ, Gönci B, Jurányi Z, Selmeczi D, Vicsek T. Phase transition in the collective migration of tissue cells: Experiment and model. Physical Review E - Statistical, Nonlinear, and Soft Matter Physics. 2006;doi:10.1103/PhysRevE.74.061908.

26. Trepat X, Wasserman MR, Angelini TE, Millet E, Weitz DA, Butler JP, et al. Physical forces during collective cell migration. Nature Physics. 2009;doi:10.1038/nphys1269.

27. Bi D, Yang X, Marchetti MC, Manning ML. Motility-driven glass and jamming transitions in biological tissues. Physical Review X. 2016;doi:10.1103/PhysRevX.6.021011.

28. Merkel M, Manning ML. Using cell deformation and motion to predict forces and collective behavior in morphogenesis; 2017.

29. Brugués A, Anon E, Conte V, Veldhuis JH, Gupta M, Colombelli J, et al. Forces driving epithelial wound healing. Nature Physics. 2014;doi:10.1038/NPHYS3040.

30. Ferrier J, Ross SM, Kanehisa J, Aubin JE. Osteoclasts and osteoblasts migrate in opposite directions in response to a constant electrical field. JCell Physiol. 1986;129(0021-9541 (Linking)):283–288.

31. Hiemer B, Ziebart J, Jonitz-Heincke A, Grunert PC, Su Y, Hansmann D, et al. Magnetically induced electrostimulation of human osteoblasts results in enhanced cell viability and osteogenic differentiation. International Journal of Molecular Medicine. 2016;38(1):57–64. doi:10.3892/ijmm.2016.2590.

32. Kaivosoja E, Sariola V, Chen Y, Konttinen YT. The effect of pulsed electromagnetic fields and dehydroepiandrosterone on viability and osteo-induction of human mesenchymal stem cells. Journal of Tissue Engineering and Regenerative Medicine. 2015;9(1):31–40. doi:10.1002/term.1612.

33. Brighton CT, Hozack WJ, Brager MD, Windsor RE, Pollack SR, Vreslovic EJ, et al. Fracture healing in the rabbit fibula when subjected to various capacitively coupled electrical fields. Journal of Orthopaedic Research. 1985;3(3):331–340. doi:10.1002/jor.1100030310.

34. Jacobs DJ, Trivedi D, David C, Yengo CM. Kinetics and thermodynamics of the rate-limiting conformational change in the actomyosin V mechanochemical cycle. J MolBiol. 2011;407(0022-2836 (Linking)):716–730.

35. Sich NM, O’Donnell TJ, Coulter SA, John OA, Carter MS, Cremo CR, et al. Effects of actin-myosin kinetics on the calcium sensitivity of regulated thin filaments. J BiolChem. 2010;285(0021-9258 (Linking)):39150–39159.

36. Stains JP, Civitelli R. Cell-to-cell interactions in bone; 2005.

37. Stains JP, Fontana F, Civitelli R. Intercellular junctions and cell-cell communication in the skeletal system. In: Principles of Bone Biology; 2019.

38. Lochner K, Fritsche A, Jonitz A, Hansmann D, Mueller P, Mueller-Hilke B, et al. The potential role of human osteoblasts for periprosthetic osteolysis following exposure to wear particles. International Journal of Molecular Medicine. 2011;doi:10.3892/ijmm.2011.778.

